# Nasty Prophages and the Dynamics of Antibiotic-Tolerant Persister Cells

**DOI:** 10.1101/200477

**Authors:** Alexander Harms, Cinzia Fino, Michael A. Sørensen, Szabolcs Semsey, Kenn Gerdes

**Affiliations:** Centre of Excellence for Bacterial Stress Response and Persistence, Department of Biology, University of Copenhagen, Copenhagen N, Denmark

## Abstract

Bacterial persisters are phenotypic variants that survive antibiotic treatment in a dormant state and can be formed by multiple pathways. We recently proposed that the second messenger (p)ppGpp drives *Escherichia coli* persister formation through protease Lon and the activation of toxin-antitoxin (TA) modules. This model found support in the field, but also generated controversy as part of recent heated debates on the validity of significant parts of the literature. In this study, we therefore used our previous work as a model to critically examine common experimental procedures in order to understand and overcome the inconsistencies often observed between results of different laboratories. Our results show that seemingly simple antibiotic killing assays are very sensitive to variation of culture conditions and bacterial growth phase. Additionally, we found that some assay conditions cause the killing of antibiotic-tolerant persisters via induction of cryptic prophages. Similarly, the inadvertent infection of mutant strains with bacteriophage φ80, a notorious laboratory contaminant, has apparently caused several phenotypes that we reported in our previous studies. We therefore reconstructed all infected mutants and probed the validity of our model of persister formation in a refined assay setup that uses robust culture conditions and unravels the dynamics of persister cells through all bacterial growth stages. Our results confirm the importance of (p)ppGpp and Lon, but do not anymore support a role of TA modules in *E. coli* persister formation. We anticipate that the results and approaches reported in our study will lay the ground for future work in the field.

**Importance:** The recalcitrance of antibiotic-tolerant persister cells is thought to cause relapsing infections and antibiotic treatment failure in various clinical setups. Previous studies have identified multiple genetic pathways involved in persister formation, but also revealed reproducibility problems that sparked controversies about adequate tools to study persister cells. In this study we unraveled how typical antibiotic killing assays often fail to capture the biology of persisters and instead give widely different results based on ill-controlled experimental parameters and artifacts caused by cryptic as well as contaminant prophages. We therefore established a new, robust assay that enabled us to follow the dynamics of persister cells through all growth stages of bacterial cultures without distortions by bacteriophages. This system also favored adequate comparisons of mutant strains with aberrant growth phenotypes. We anticipate that our results will contribute to a robust, common basis of future studies on the formation and eradication of antibiotic-tolerant persisters.

## Introduction

Bacterial persisters constitute a subpopulation of phenotypically antibiotic-tolerant cells forming among a population of genetically antibiotic-susceptible bacteria. Persister cells are usually slow-or nongrowing, and the field largely agrees that the antibiotic tolerance of persisters is linked to a dormant physiological state in which the cellular processes commonly poisoned by bactericidal antibiotics are inactive (1, 2). Consistently, a large body of studies from several laboratories has uncovered genetic pathways that control and execute the formation of persister cells of *Escherichia coli* K-12 as a phenotypic conversion into dormancy (1, 2). These mechanisms include a drop of cellular ATP levels, the modulation of nucleoid-associated proteins, changes in metabolic fluxes, high expression of drug efflux pumps, or the activation of different sets of toxin-antitoxin (TA) modules (3–10). The emerging picture is therefore that various parallel and partially interlinked pathways of persister formation in *E. coli* K-12 give rise to a heterogeneous population of persisters that have formed through different pathways, have different physiological properties, and thus exhibit different profiles of antibiotic tolerance (1, 2, 11).

The complexity of bacterial persister formation and the sensitivity of persistence assays to even slight experimental variation have generated controversies that made the field notorious for debates on technical and biological aspects of studying bacterial persistence (12–16). Among the work cited above, our laboratory has published a series of studies linking persister formation in *E. coli* K-12 to the activation of a set of ten mRNA endonuclease toxin TA modules under control of the second messenger (p)ppGpp, polyphosphate, and protease Lon (8, 9). Though similar findings have been made, e.g., in *Salmonella* Typhimurium (17), our model has also been met with skepticism by other researchers in the field (5, 13, 14, 18). In the present study we therefore carefully re-evaluated our previous conclusions as well as the underlying methodology and also probed common experimental procedures for sources of the inconsistencies frequently observed between studies in the field.

In short, we discovered that *E. coli* mutant strains used in our previous work had been inadvertently infected with several different lysogenic bacteriophages and that prophage carriage strongly affected persistence measurements. However, we also show that already the resident cryptic prophages of the *E. coli* K-12 MG1655 wild type strain distort the results of persister assays under conditions that are commonly used in the field. Importantly, we show that the common practice of determining persister levels at only a single growth time-point is often inappropriate to account for shifted dynamics of growth and persister formation in mutant strains. We finally tested the key components of our previously proposed model of persister formation using new mutant strains and a refined methodology. Crucially, we could confirm a role of (p)ppGpp, polyphosphate, and Lon in bacterial persister formation and / or survival, but did not find strong evidence for the involvement of TA modules or the connection of these components in a single pathway of persister formation as we had proposed earlier.

## Results

### Classical persister assays suffer from technical and biological drawbacks

The formation of persister cells is typically measured by determining the fraction of antibiotic-tolerant cells in bacterial cultures that are considered to be exponentially growing some hours after inoculation from dense overnight cultures (Figure 1A). A biphasic kinetic of antibiotic killing reveals the presence of persister cells, because these are killed slowly and can be detected after the regular cells have been rapidly eliminated in a first phase of killing (19). Despite the apparent simplicity of this experimental setup, persister assays are known to be sensitive to even minor variation of the experimental conditions and have often given inconsistent results in different laboratories (5, 16, 20, 21). We therefore suspected that this simple assay setup may be inadequate to represent the dynamic nature of bacterial persistence and could give results that are strongly affected by biological or technical parameters that are usually not controlled in persistence assays.

**Figure 1.**
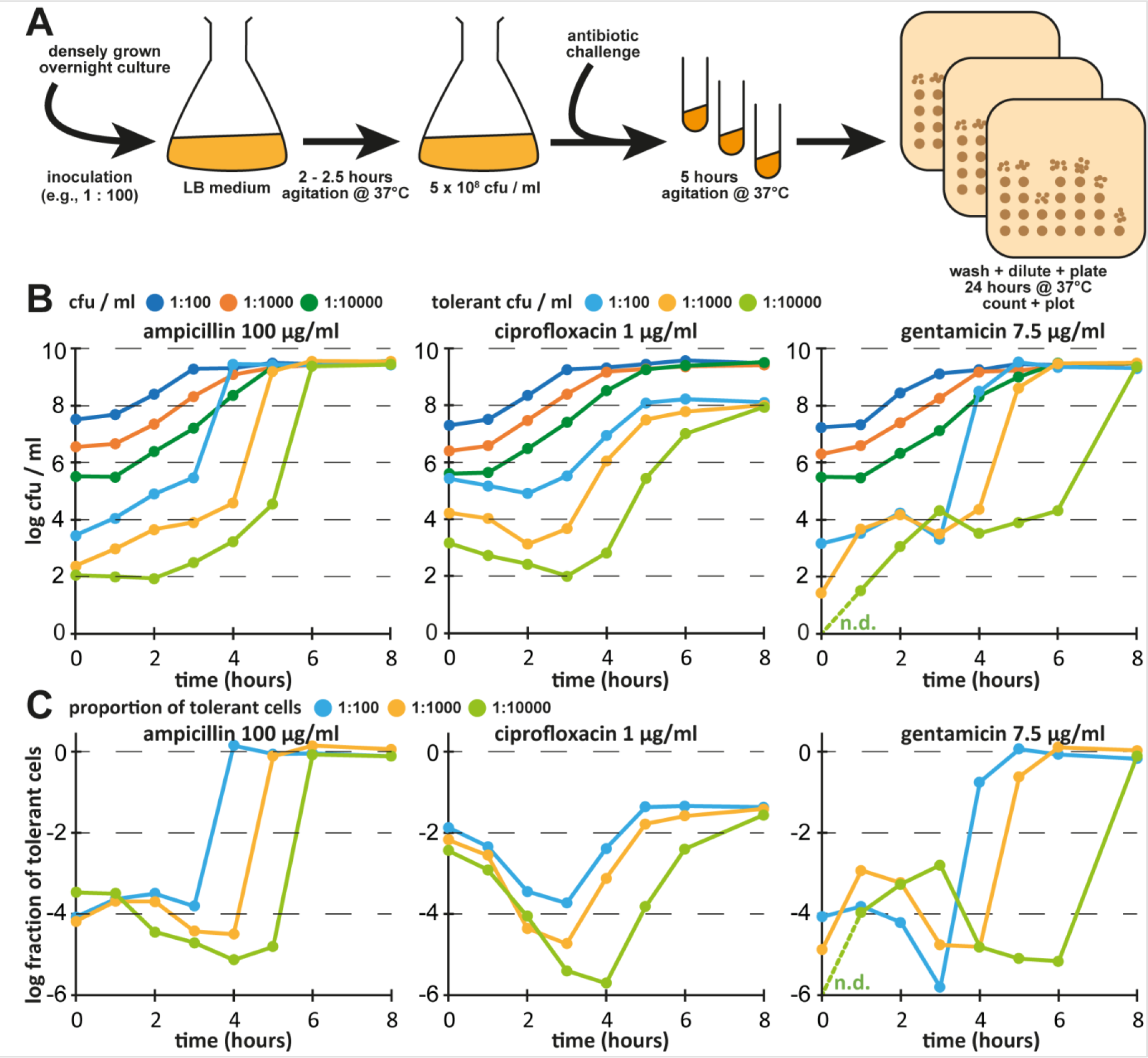
Persister assays are affected by inoculum and growth phase. (A) Scheme illustrating the setup of a persister assay as it is commonly performed in the field. (B) Cultures of *E. coli* K-12 MG1655 were grown in LB medium after inoculation 1:100 (blue), 1:1000 (orange), or 1:10000 (green) from dense overnight cultures and all colony forming units (cfu) as well as antibiotic-tolerant cfu were determined at each time-point. (C) Fraction of antibiotic-tolerant cells for each data point of (B). Data points are shown for one representative experiment, because the absolute numbers of antibiotic-tolerant cells (but not their dynamics) were affected considerably by batch-to-batch variation of the LB medium (see Figure S1). n.d. = not detected (no gentamicin-tolerant bacteria recovered at t = 0 h from inoculation 1:10000)

As a first step towards understanding the notorious variability of results reported for single growth time-point persister assays, we followed the absolute levels of colony forming units (cfu) and antibiotic-tolerant cells over time from inoculation of an *E. coli* culture through exponential growth into stationary phase in LB medium (Figures 1B and 1C). Cultures were treated with lethal concentrations of ampicillin, ciprofloxacin, or gentamicin (representing β-lactams, fluoroquinolones, and aminoglycosides that kill by very different mechanisms) for five hours that are more than sufficient to kill all regular cells at least during exponential growth so that only persisters remain (5, 8, 9, 11).

Most importantly, we found that cultures inoculated with different numbers of cells from overnight cultures displayed systematically different fractions of antibiotic-tolerant cells at any given total cfu / ml during exponential growth (e.g., at ca. 5 × 10^8^ cfu / ml that are reached after two, three, and four hours of growth when inoculated at 1:100, 1:1000, and 1:10000 dilutions from stationary phase overnight cultures, respectively; Figures 1B and 1C). As an example, the fraction of cells tolerant to 1 μg/ml of ciprofloxacin, a commonly used treatment setup, seemed to be 10^-4^, 10^-5^, or 10^-6^ at this growth stage (5 × 10^8^ cfu / ml) depending on the inoculum. This finding is incompatible with the idea that this setup of persister assays is measuring stochastic persister formation of exponentially growing bacteria, since this should be independent of the inoculum. Furthermore, the levels of tolerant cells differed massively between the three antibiotics and over time even at periods when the overall cfu/ml changed only marginally, highlighting that the heterogeneity of persister cells makes it inappropriate to simply report “persister levels” from a single time-point or a given cell density.

In addition, we observed a variation of results obtained with different batches of LB medium in a way that bacterial growth and the overall dynamics of antibiotic tolerance were not affected, but the absolute levels of tolerant cells varied considerably (Figure S1). It is well known that LB medium is prone to batch-to-batch variation that can essentially not be controlled in the laboratory, e.g., due to the degradation of L-tryptophan over time depending on the exposure to ambient light or the degree of L-asparagine and L-glutamine deamidation during autoclaving (22). Furthermore, the poor content of divalent cations and sugars in LB medium causes variation in the availability of these important nutrients to *E. coli* cultures due to even minor differences in handling during the preparation of each batch (23).

### Following the dynamics of *E. coli* persister cells in M9 medium

To overcome the issues outlined above, we decided to switch the culture medium from LB to more defined M9 medium (see details in *Materials and Methods*) and adopted the determination of antibiotic tolerance in bacterial cultures over time as our standard procedure (Figure 2A). When we studied the dynamics of *E. coli* K-12 wildtype cells tolerant to ampicillin, ciprofloxacin, or gentamicin as described above, the results of experimental replicates were much more homogeneous and showed similarities but also differences to previous experiments in LB medium (Figures 2B and 2C). For ampicillin, the level of tolerant cells remained constant after inoculation throughout exponential growth and then sharply increased during late-exponential growth until all bacteria were ampicillin tolerant in stationary phase, because this antibiotic is unable to kill non-growing cells (24). Similarly, the number of ciprofloxacin-tolerant cells was stable for the first hours and then increased into stationary phase, while – unlike with ampicillin – only up to 10% of the population became tolerant. For gentamicin, the dynamics of antibiotic-tolerant cells looked very different from those of the other antibiotics, and the levels of tolerant cells initially increased until mid-exponential phase, decreased again until entry into stationary phase, and finally rose up to almost full tolerance at the end of the experiment. We interpret these results as indicating that the stable levels of ampicillin-and ciprofloxacin-tolerant cells after inoculation and throughout early exponential growth represent dormant cells that have been carried over from the overnight cultures and not a steady state of persister formation and resuscitation. This interpretation is also in line with the direct correlation of inoculum and the fraction of antibiotic-tolerant cells that we had observed before (Figures 1B and 1C).

**Figure 2.**
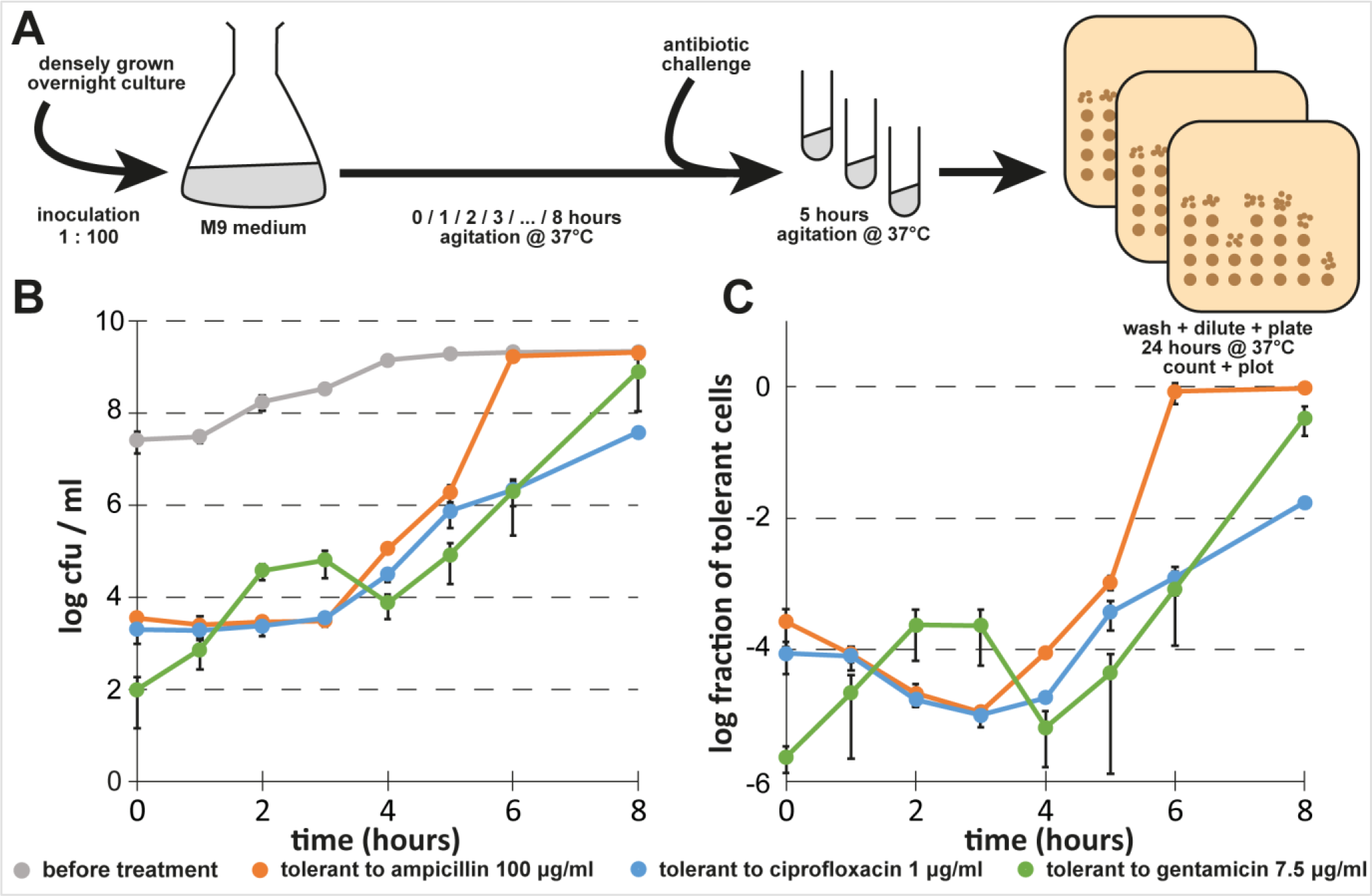
Persister formation of *E. coli* K-12 MG1655 in M9 medium. (A) Scheme illustrating how the dynamics of persister formation were determined. (B) Bacterial growth and the dynamics of antibiotic-tolerant cells were determined for cultures of *E. coli* K-12 MG1655 in M9 minimal medium as described for the experiment shown in Figure 1 after inoculation 1:100 from a dense overnight culture. (C) Fraction of antibiotic-tolerant cells for each data point shown in (B). We generated biphasic kill curves to verify that antibiotic-tolerant cells during exponential growth (3 hours after inoculation) are persisters (Figure S2). Data points represent the mean of at least three independent experiments and error bars indicate the standard deviation.

### Persister assays with ciprofloxacin in *E. coli* K-12 are affected by induction of resident prophages

Beyond our worries regarding the growth conditions of bacteria studied in persistence assays, a recent publication of the Brynildsen laboratory on *Staphylococcus aureus* persister formation alerted us that the presence of resident prophages may affect measurements of persister frequencies under some conditions (25). In short, these authors showed that ciprofloxacin treatment at commonly used concentrations of 0.5-1 μg/ml – more than an order of magnitude above the minimum inhibitory concentration (MIC) and sufficient to kill all regular cells – eliminates a substantial fraction of persisters not due to the direct action of the antibiotic, but due to the secondary activation of prophages. This effect could be overcome at very high concentrations of ciprofloxacin that cause a strong inhibition of cellular DNA processing and thus impair prophage development (25). Given that DNA damage is generally known to be a strong inducer of temperate prophages including some of the cryptic prophages in the *E. coli* K-12 MG1655 chromosome (26, 27), we suspected that a similar effect as described by E. L. Sandvik, et al. (25) may also affect persister results reported for *E. coli*. We therefore compared *E. coli* K-12 wild type and a previously published mutant devoid of all nine cryptic prophages for tolerance to different concentrations of ciprofloxacin in a classical single growth time-point persister assay (Figure 3). Similar to the results reported for *S. aureus*, the *E. coli* K-12 wild type showed a minimum of survival at commonly used intermediate concentrations of ciprofloxacin (0.5-1 μg/ml) and displayed greatly increased survival at higher concentrations (Figure 3A). This effect was abolished in the mutant lacking the cryptic prophages (Figure 3B). When we studied the gap between survival of 1 μg/ml and 10 μg/ml ciprofloxacin treatment over time, we found that the difference was considerable (around 1 log) throughout all growth phases but most pronounced during exponential growth (Figures 3C and 3D). In order to avoid artifacts from the induction of cryptic prophages, we therefore adopted 10 μg/ml as the standard concentration of ciprofloxacin for antibiotic killing assays.

**Figure 3.**
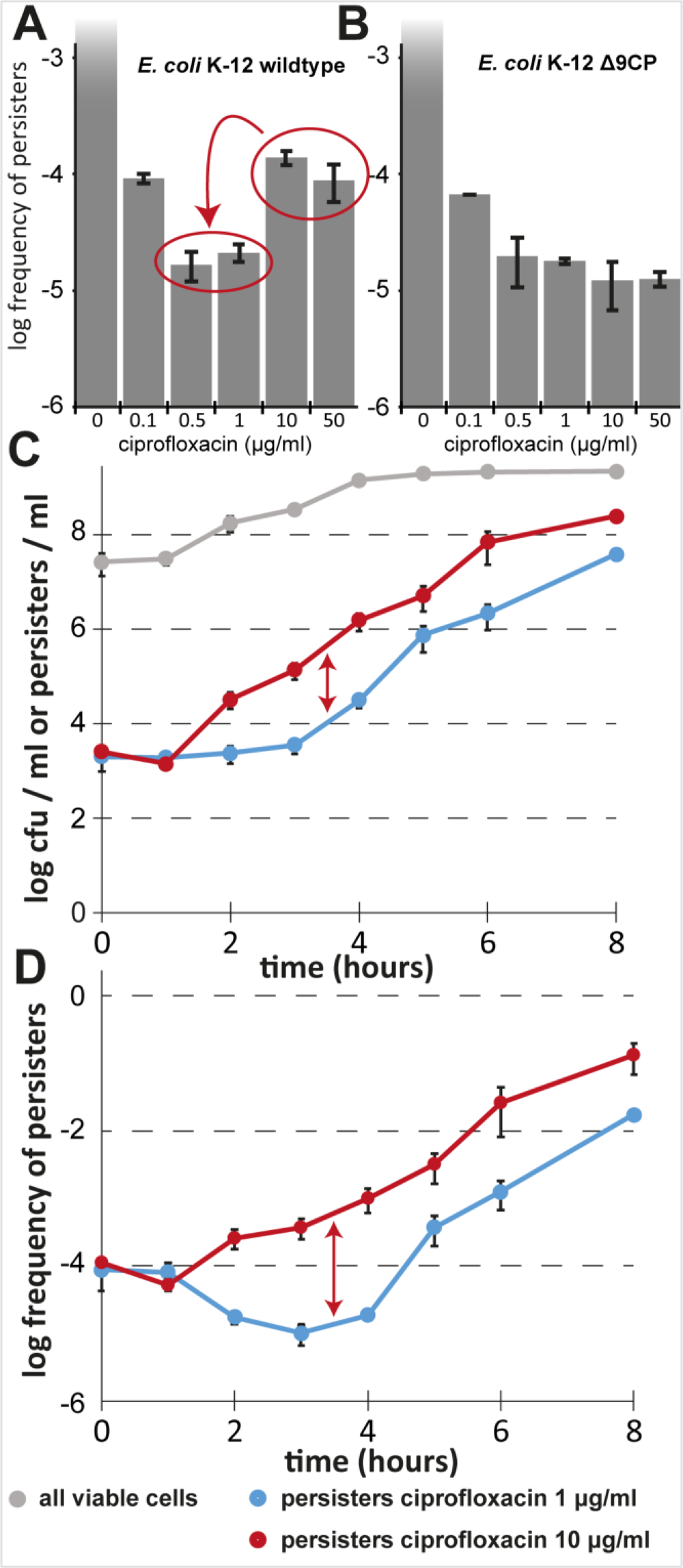
Induction of cryptic prophages distorts persister measurements of *E. coli* K-12. Exponentially growing cultures of *E. coli* K-12 strain BW25113 (A) and its derivative lacking all nine cryptic prophages (B; Δ*9CP*) of X. Wang, et al. (27) were treated with different concentrations of ciprofloxacin in M9 medium and the level of surviving persister cells was determined. Note that the level of survivors for the wildtype strain increases at high ciprofloxacin concentrations, while no such effect can be observed in the Δ9CP mutant. The latter strain generally exhibits a lower level of persister formation than its ancestor, possibly due to the roles of prophage-encoded factors in persistence and general stress tolerance (27). (C) Bacterial growth and the dynamics of antibiotic-tolerant cells were determined for cultures of *E. coli* K-12 MG1655 in M9 minimal medium as described for Figure 2B and the treatment with 1 μg/ml and 10 μg/ml ciprofloxacin were compared. The fraction of antibiotic-tolerant cells at each time-point is shown in (D). Data points represent the mean of at least three independent experiments and error bars indicate the standard deviation.

### *E. coli Δ1-10TA* carry a defective lambda prophage

The major effect of cryptic prophages on the detection of ciprofloxacin-tolerant persisters caught our attention, because a previous analysis of the genome sequence of our *E. coli* K-12 MG1655 *Δ10TA* strain by Kim Lewis’ laboratory had revealed an accumulation of nucleotide polymorphisms in the cryptic prophages (5). The *Δ10TA* strain had been created by the sequential deletion of ten mRNA endonuclease toxin TA modules and was described to display a strong defect in bacterial persistence that has been a cornerstone of our previous work (8, 9). Worryingly, we recently discovered that the published variant of *E. coli Δ10TA* carried a lambda prophage. The infection was easily cured genetically (see *Materials and Methods*), and the resulting strain, *E. coli* K-12 MG1655 *Δ10TA attB(+)*, did not show any difference to parental *E. coli* K-12 MG1655 *Δ10TA λ(+)* in persister levels (see below). However, a general worry about prophage-mediated effects on persistence and the sustained debate over the results of our previous work prompted us to sequence the genome of *E. coli* K-12 MG1655 *Δ10TA attB(+)*. Additionally, we sequenced two ancestors of *Δ10TA*, *Δ5TA* and *Δ8TA*, in which five and eight mRNA endonuclease toxin TA modules had been deleted, respectively (9). Notably, *Δ5TA* had been the first strain in the series of sequential TA module deletions of our previous work that was found to exhibit a defect in persister formation (9).

Surprisingly, the genome sequences showed that *Δ5TA* and *Δ8TA* carried a lambda prophage at the *attB* site in the *gal-bio* region of the *E. coli* chromosome (Figure 4A). We therefore tested the most important strains of our previous studies for lambda infection by PCR and found that every TA module deletion strain of the series leading to *Δ10TA* as well as the *relA spoT* mutant (deficient in (p)ppGpp signaling) were lambda lysogens (Figure 4B). Interestingly, the genome sequences of *Δ5TA* and *Δ8TA* revealed a deletion of ca. 13 kb inside the lambda prophage around the major repressor genes *cl, cll*, and *clll* and major promoters, suggesting that the prophage is defective and largely inactive (Figure 4A).

**Figure 4.**
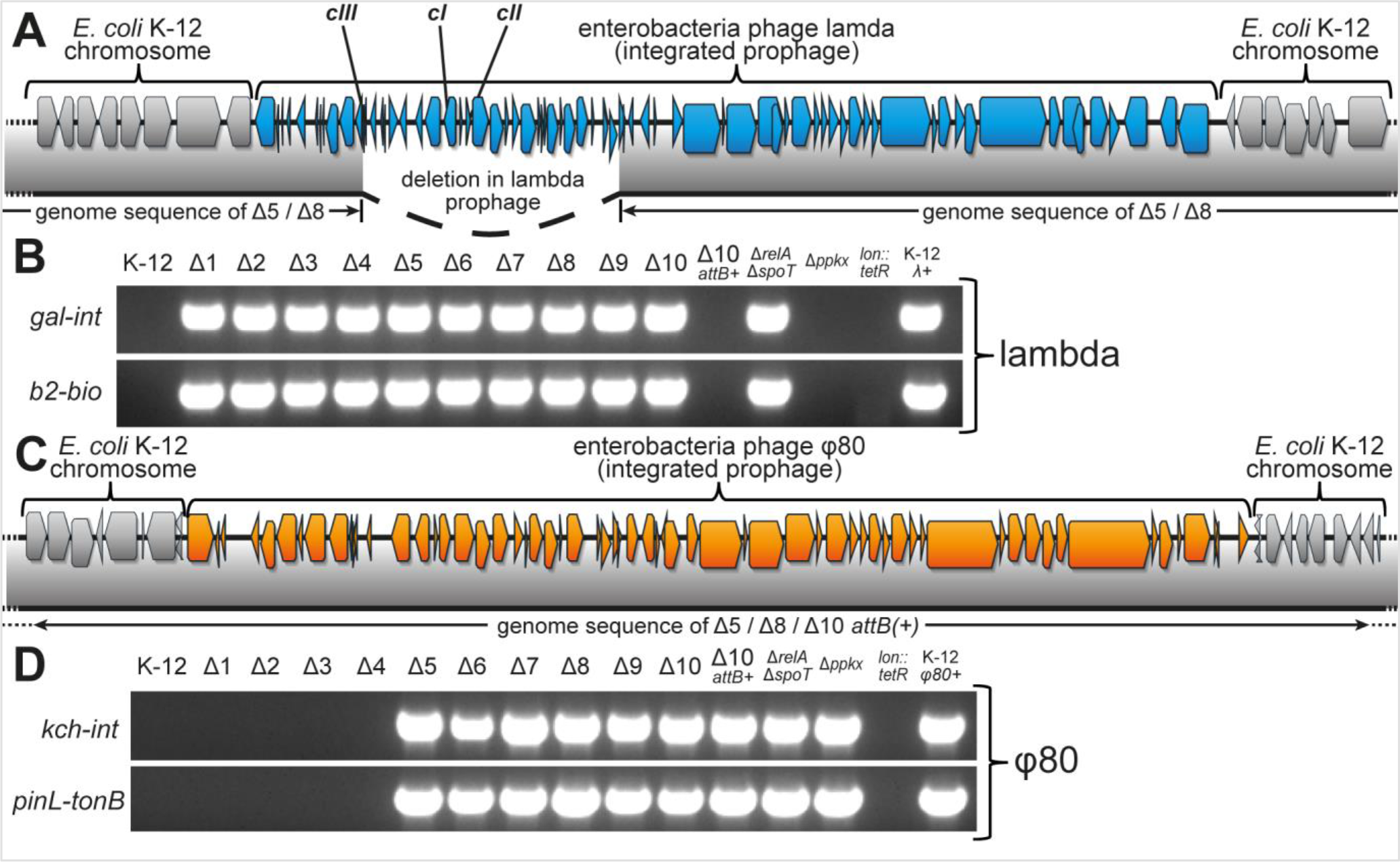
Lambda and φ80 infection of strains from our previous studies. (A) The insertion of a lambda prophage (blue gene arrows) in the *E. coli* K-12 chromosome (grey gene arrows) is shown together with coverage of the prophage insertion in the genomes of Δ*5TA* and Δ*8TA* (grey bar below). The 13 kilobase deletion comprising *cl, cll,* and *clll* repressors spans from the nuclease inhibitor gene *gam* to the endolysin gene R. (B) Diagnostic PCRs over the two junctions between lambda prophage and the gal-bio region of the *E. coli* chromosome were performed to determine the extent of lambda infection in important strains from our previous studies (8, 9). (C) Insertion of a φ80 prophage (grey gene arrows) in the *E. coli* K-12 chromosome (grey gene arrows) in the genomes of Δ*5TA*/Δ*8TA*/Δ*10TA attB(+)*. (D) Diagnostic PCRs over the two junctions between the φ80 prophage and the *ycil* locus in the *E. coli* K-12 chromosome were performed to determine the prevalence of φ80 infections among important strains from our previous work (8, 9).

### Widespread infection with φ80 in strains used to support the model

Beyond a defective lambda prophage in *Δ5TA* and *Δ8TA*, we discovered that all three sequenced strains of our TA module deletion series carried a φ80 prophage at its *attP* integration site in the *ycil* gene (Figure 4C). We therefore performed diagnostic PCRs on the junctions of φ80 integration in all strains that were crucial for the main results of our previous articles (8, 9). Worryingly, these experiments revealed φ80 infections in *Δ5TA-Δ10TA, relA spoT*, and *ppkx* (polyphosphate metabolism) mutants (Figure 4D).

In parallel to studying the φ80 infections, we investigated whether the lambda prophages in our strains all carried the 13 kb deletion revealed for *Δ5TA* and *Δ8TA* by testing their susceptibility to the lambda *cl*_*b221*_ mutant encoding an inactive *cl* repressor gene. The lambda *cl*_*b221*_ mutant is obligatory lytic unless *cl* is provided *in trans* e.g., by a lambda prophage. Surprisingly, the *Δ10TA attB(+)* strain was immune to lambda *cl*_*b221*_ infection although it had been cured of the lambda prophage, but all TA module deletion strains before and including *Δ8TA* were sensitive (Figure 5A). These results show that the lambda prophage had already been inactivated in *Δ1TA* and may already have been present in the ancestral *E. coli* K-12 MG1655 wildtype stock. The surprising immunity of *Δ9TA* and *Δ10TA* to lambda *cl*_*b221*_ suggested that these strains must encode a source of lambda *cl* outside of the defective prophage that lacks *cl*. Consistently, a deeper analysis of the *Δ10TA attB(+)* genome sequence identified a 12 kb piece of lambda centered around the *cl* gene (Figure 5B). Interestingly, the flanking regions of this lambda segment were φ80 sequences, not part of the *E. coli* K-12 MG1655 genome, indicating that these strains had been infected with a φ80/lambda hybrid on top of the φ80 wildtype phage. Given that the genetic architecture of these two phages is very similar, hybrid phages are often viable and had been used as tools to study different features of phage biology in earlier times (28). A comparison of the *Δ10TA attB(+)* genome sequence with published φ80/lambda hybrids revealed that the hybrid phage in our strains is known as φ80_*h(80)imm(λ)*_, a phage that produces φ80 particles but carries a lambda immunity region (Figure 5B, (28)).

**Figure 5.**
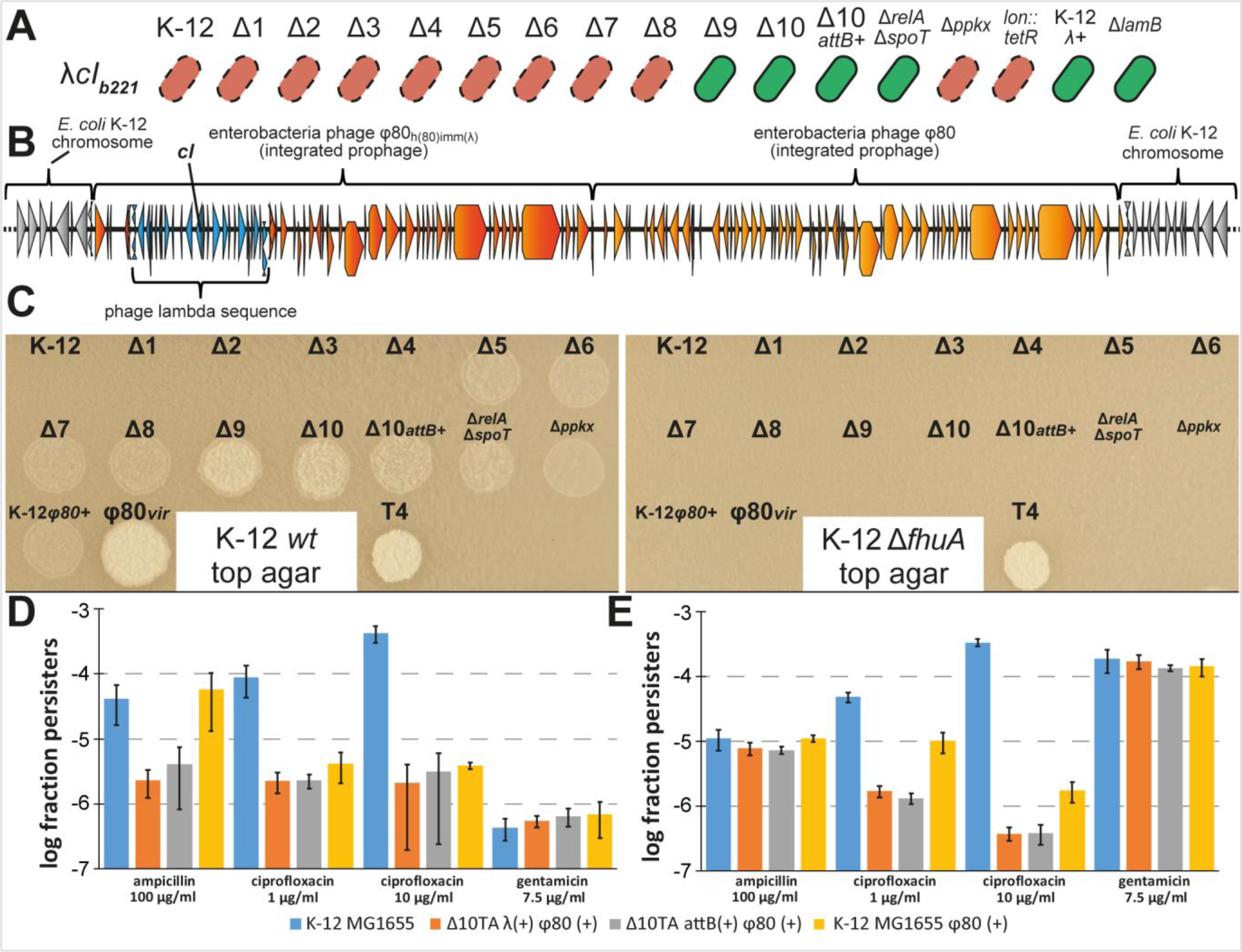
Lambda immunity, active φ80 prophages, and their effect on persistence. (A) The sensitivity of different *E. coli* strains to the lambda *cl*_*b221*_ mutant was determined by streaking across lines of phage stock on agar plates. red = no growth / sensitive, green = growth / immune (B) Illustration of neighboring φ80 and φ80_*h(80)imm(λ)*_ integration in the chromosome of *E. coli* Δ*10TA* as deduced from the genome sequence. The *E. coli* chromosome backbone is shown in grey, genes of φ80 and φ80_*h(80)imm(λ)*_ in orange and dark orange, respectively, and the lambda region of φ80_*h(80)imm(λ)*_ is highlighted in blue. Note that the genome sequence does not allow to determine which of the prophages is upstream and downstream in the integration site. (C) Plaque assay with culture supernatants and control phage stocks to determine the infectivity of prophage carriers. (D, E) Exponentially growing *E. coli* K-12 MG1655 and different mutant derivatives / lysogens were challenged with ampicillin, ciprofloxacin, or gentamicin in LB medium (D) or M9 medium (E) and the fractions of surviving persisters were calculated. Data points are averages of three independent experiments and error bars represent the standard deviation.

Unlike the lambda prophage, the φ80 prophages seemed complete and thus likely active. This observation was worrying, because φ80 is known as a highly infectious laboratory contaminant (29, 30) and we had widely distributed the strains of our TA module deletion series to other research groups. We therefore performed plaque assays on regular overnight cultures of various strains in order to determine the infectivity of φ80 present in our lysogens. As expected, culture supernatants of all strains that had been found to carry a φ80 prophage by PCR (Figure 4C) formed plaques on a lawn of *E. coli* K-12 wildtype cells, but not on a *fhuA* mutant that lacks the φ80 receptor (Figure 5C).

### Lysogenization with φ80 reduces persister levels similar to TA module deletions in *Δ10TA*

We were particularly worried about the φ80 lysogenization because this prophage is known to be easily induced by DNA damage (29) and this property might reduce the number of survivors of ciprofloxacin treatment, the major antibiotic that was used in our previous study (8). We therefore lysogenized the *E. coli* K-12 MG1655 wildtype using φ80 particles from the supernatant of *Δ10TA attB(+)* and assayed persister formation of this strain in direct comparison to the parental wildtype and the *Δ10TA* mutants (Figure 5D). When the experiment was performed as a single growth time-point assay in LB medium like in our previous work, we readily reproduced the significant drop of ampicillin and ciprofloxacin survival that had been reported for the *Δ10TA* mutants compared to *E. coli* K-12 wildtype (8, 9). Lysogenization of *E. coli* K-12 with φ80 had no effect on ampicillin tolerance but caused the same drop of ciprofloxacin survival that had been a leading phenotype of *Δ10TA* and other infected strains in our previous studies (Figure 5D; 8, 9). Furthermore, the difference between *E. coli* K-12 wildtype and the *Δ10TA* mutants in ampicillin tolerance disappeared when the experiment was performed in M9 minimal medium (Figure 5E), as had already been reported by Y. Shan, et al. (5). Taken together, these results directly questioned whether there was any φ80-independent difference between *E. coli* K-12 wildtype and the *Δ10TA* mutants in persister formation or survival. Interestingly, in M9 minimal medium the *Δ10TA* strains displayed clearly reduced survival of ciprofloxacin treatment when compared to the φ80 lysogen, possibly due to some of the additional mutations present in the *Δ10TA* genome revealed by Y. Shan, et al. (5) or due to the additional presence of φ80_*h(80)imm(λ)*_ Unlike what we had observed for the cryptic prophages of *E. coli* K-12 (Figure 3), an increase in the ciprofloxacin concentration from commonly used 1 Mg/ml to 10 Mg/ml did not reduce the drop in survival caused by lysogenization with φ80 (Figures 5D and 5E). It seems likely that this different behavior is due to the high sensitivity of φ80 induction to even slight DNA damage that may remain after the end of ciprofloxacin treatment and persister resuscitation (29), while the induction of cryptic prophages of *E. coli* K-12 requires very strong DNA damage (27). No differences between *Δ10TA* with and without the defective lambda prophage in the *attB* site or between any strains regarding gentamicin tolerance could be detected (Figures 5D and 5E).

### (p)ppGpp, Lon, and polyphosphate in *E. coli* K-12 persister formation

The ability of φ80 carriage to cause the same drop in persistence that we had previously attributed to the deletion of ten TA modules was worrying, because we had found this prophage in nearly all mutant strains that supported the model of persister formation proposed by our previous work (8). We therefore constructed new, uninfected versions of all mutants that had been found to be φ80 lysogens and assayed the dynamics of persister formation of these strains in order to determine whether the model still holds (see Figure 6A for a summary of the model and *Materials and Methods* for strain construction).

This model is based on the second messenger (p)ppGpp that plays important roles in persister formation of diverse organisms (2, 31). Consistently, we had previously reported severe defects in the formation or survival of ampicillin-and ciprofloxacin-tolerant persisters for mutants deficient in (p)ppGpp synthesis (*relA spoT* strains; (8)). Subsequently, there had been some debate in the field whether pleiotropic phenotypes of these mutants like slow growth, long lag times, and reduced stationary phase viability might have affected our results (5, 16). We therefore compared the dynamics of antibiotic tolerance in cultures of a *relA spoT* mutant of *E. coli* K-12 MG1655 to the parental wildtype in a variant of our M9 medium that had been supplemented with a mixture of amino acids to support growth at roughly the same rate for both strains (see *Materials and Methods* as well as K. Potrykus, et al. (32)). Under these conditions, we observed a considerable defect of the *relA spoT* mutant in tolerance to ciprofloxacin and gentamicin throughout all growth phases, though the overall shape of the curves following the levels of tolerant cells over time was only poorly affected (Figures 6B and 6C). The *relA spoT* mutant also seemed to display a defect in tolerance to ampicillin during exponential growth, though the very different final cell densities and the early onset of full ampicillin tolerance upon cessation of growth by the *relA spoT* mutant made it difficult to judge this phenotype (Figures 6B and 6C).

**Figure 6.**
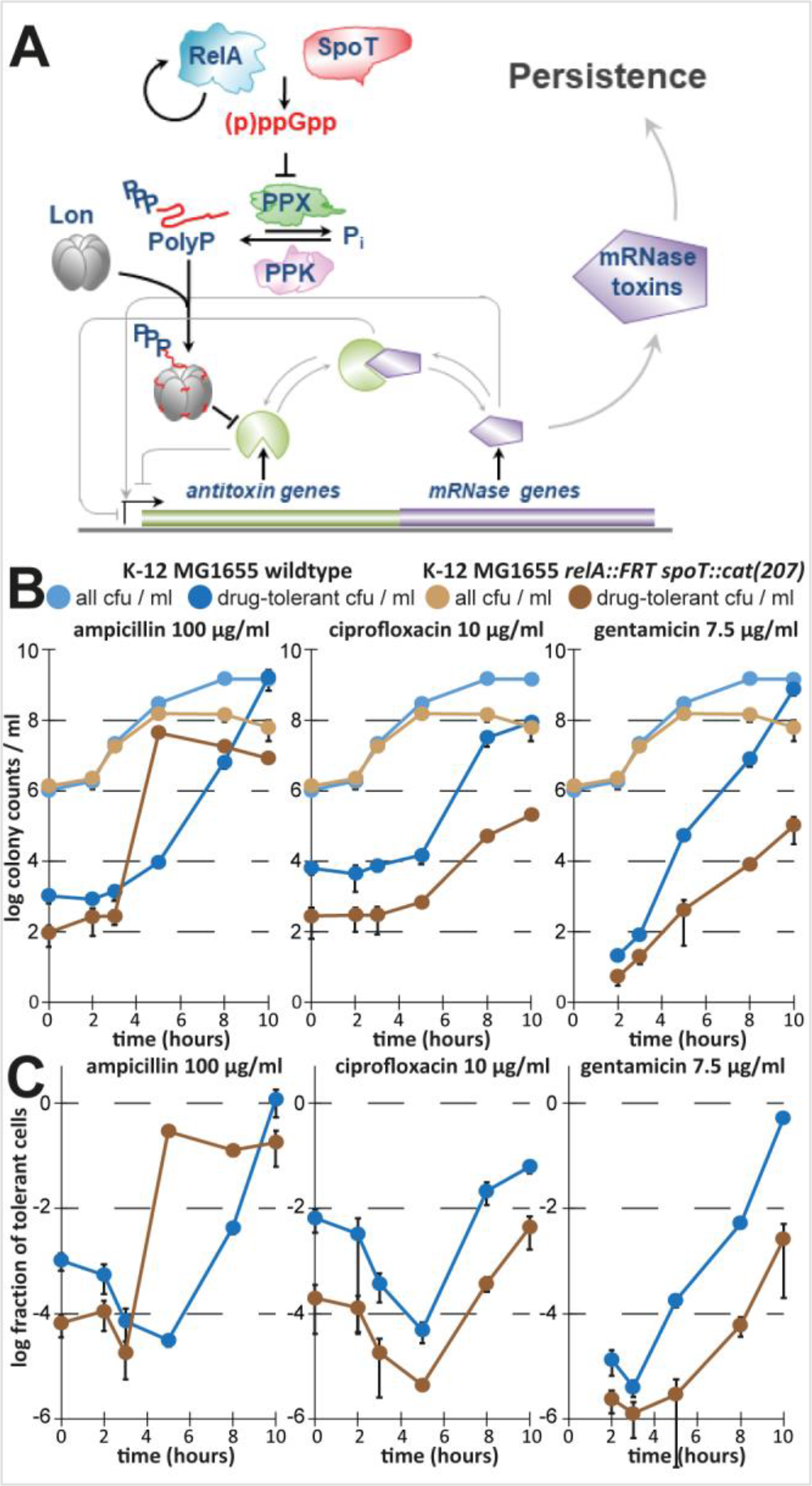
The model of (p)ppGpp-dependent persister formation through TA modules. (A) The illustration (adapted from E. Germain, et al. (53)) shows our previously published model of persister formation initiated by stochastic bursts of (p)ppGpp that induce the production of polyphosphate which stimulates Lon to degrade TA module antitoxins. Consequently, the activation of mRNA interferase toxins would induce bacterial persistence (8). (B, C) In order to verify the most upstream element of the model, we created a new (p)ppGpp-deficient mutant of *E. coli* K-12 MG1655 (*relA spoT*) and assayed the dynamics of antibiotic tolerance as described for the experiment in Figure 2 with minor modifications (see *Materials and Methods*).

Downstream of (p)ppGpp, the model proposed by E. Maisonneuve, et al. (8) comprises the production of polyphosphate by PPK and degradation by PPX (both lacking in *Δppkx* that produces little polyphosphate), the polyphosphate-dependent activation of Lon (impaired in a *lon* deletion), and the degradation of TA module antitoxins by Lon to activate toxins and induce persistence (impaired in a *Δ10TA* knockout; Figure 6A). Notably, the only of these mutant strains of our previous studies that showed a defect in bacterial persistence but had not been infected with φ80 was the *lon::tet* strain (Figure 4D; (8, 9)). However, the use of *lon* single mutants is prone to artifacts in persister assays due to the activation of SulA, an inhibitor of cell division, in response to DNA damage which is essentially irreversible in the absence of Lon that degrades SulA (26, 33). We therefore created a *sulA lon* double mutant similar to the one that was used by A. Theodore, et al. (33) in order to exclude these artifacts. This mutant showed a defect in the formation or survival of ciprofloxacin-tolerant persisters during exponential growth and also exhibited a markedly slower increase in ampicillin tolerance during stationary phase (Figure 7). Conversely, the *ppkx* mutant showed a drop of ciprofloxacin tolerance only in stationary phase and was, beyond a slightly shifted curve likely caused by its slower growth, not affected in ampicillin tolerance (Figure 7). We therefore conclude that our results confirm roles of (p)ppGpp as well as the Lon protease in the formation of persister cells by *E. coli* K-12, but these data cannot verify whether they are part of the same pathway as suggested previously or not. Furthermore, we failed to detect any phenotype of a newly constructed *Δ10TA* strain (called *Δ10’TA*) without prophages in tolerance to any of the tested antibiotics at any time-point when compared to the parental wildtype strain (Figure 7). It appears therefore that unnoticed lysogenization with φ80 has been the reason for all persister phenotypes that we detected with the initial *Δ10TA* strain.

**Figure 7.**
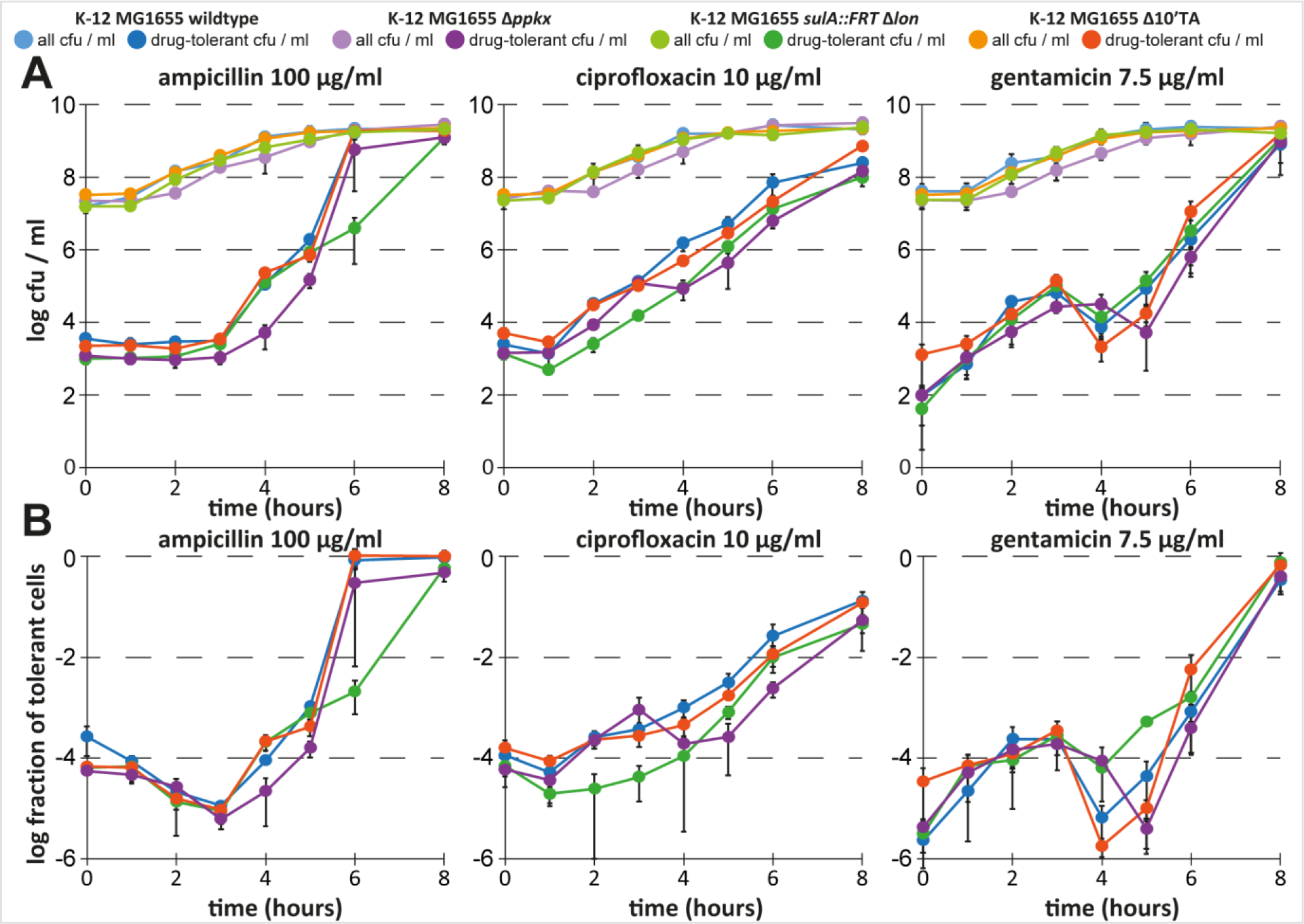
Protease Lon, but not mRNA endonuclease TA modules, contribute to persistence. We studied the dynamics of antibiotic tolerance for cultures of newly constructed *E. coli* K-12 Δ*ppk×*, *sulA::FRT Δlon*, and *Δ10’TA* mutants in comparison to the parental wildtype similar to the experiment shown in Figure 2. Changes in overall and antibiotic-tolerant cfu/ml are plotted in (A) and the fractions of antibiotic-tolerant cells are plotted in (B). While the *sulA::FRT Δlon* mutant shows a clear defect in persister formation or survival during exponential growth (around 3 hours after inoculation; see also Figure S2), the parental *sulA::FRT* strain had no phenotype (Figure S3), confirming that the defect is due to the lack of Lon.

## Discussion

### Persister phenotypes as an “inside job” of φ80

In this study we showed that important aspects of our previously published model of persister formation were based on the misinterpretation of experimental results caused by the unnoticed lysogenization of many of our strains with bacteriophage φ80. Consequently, defects in persister formation that we had previously attributed to different genetic mutations are instead a consequence of φ80 prophages residing in the chromosomes of these strains.

Others had previously questioned our model of persister formation based on a number of different considerations. Most importantly, some studies did not observe phenotypes of strains deficient in important components of our model in single growth time-point persister assays under their conditions. For example, Y. Shan, et al. (5) did not detect a defect of *lon sulA* and *ppkx* strains in persister formation or survival upon treatments with ampicillin and ciprofloxacin. Also, while the role of (p)ppGpp as an important controller of persister formation is generally well established (2, 31), recent studies suggested that the link between (p)ppGpp signaling and the activation of TA modules needs to be reinvestigated. As an example, a dedicated article by B. C. Ramisetty, et al. (18) and an associated commentary by L. Van Melderen and T. K. Wood (13) raised the point that some of the antitoxins of the ten TA modules implicated in our model may not be targets of the Lon protease. Furthermore, they used the *yefM-yoeB* TA module as a model to study the link between (p)ppGpp and TA module activation and reported that polyphosphate was, in their hands, not required for *yefM-yoeB* activation after experimental induction of (p)ppGpp signaling or Lon overexpression (18). We agree that the role of polyphosphate requires further investigation and, with the refined methodology of our current study, we could not confirm a role of this molecule in the formation and survival of persister cells during exponential growth of *E. coli* K-12 (Figure 7). For Lon, others had previously argued that our failure to use a *sulA lon* double mutant might be the reason why we detected persister phenotypes of Lon-deficient strains that they were unable to reproduce (5, 33). In our hands, the *sulA lon* double mutant shows a clear 1-log reduction of ciprofloxacin-tolerant persisters during exponential growth (Figure 7), supporting our previous observations (8) and suggesting that the degradation of Lon targets – either specific targets like antitoxins or simply unfolded proteins – is important for persister formation or survival. Future studies should therefore further investigate the links between (p)ppGpp, protease Lon, and bacterial persistence.

Beyond Lon and polyphosphate, considerable criticism revolved around the question whether our *Δ10TA* strain was a useful tool to study the link between TA modules and bacterial persistence. Others have reported various signs of slightly decreased viability of the original *Δ10TA* strain like a minor drop in MIC to several antibiotics that they traced back to polar effects of our TA module deletions or additional mutations acquired during the long series of recombineering steps (13, 18). Notably, changes in MIC should not affect persistence *per se* (19), but it is entirely possible that these phenotypes of *Δ10TA* are caused by the φ80 prophage that indeed increases the sensitivity of *E. coli* K-12 to, e.g., DNA-damaging agents (29). Beyond ampicillin and ciprofloxacin, we have in this study systematically included the aminoglycoside gentamicin in our antibiotic killing assays but did (apart from the *relA spoT* mutant; Figure 6) not detect any relevant differences in tolerance to this antibiotic. We chose gentamicin as a control because others showed previously that persister cells formed through the activity of mRNA endonuclease toxins should not be appreciably tolerant to aminoglycoside antibiotics, since these toxins fail to fully shut down translation and thus cannot prevent its poisoning by aminoglycoside drugs (34). Notably, other mechanisms of persister formation like the activation of different TA modules (with small peptide toxins that abrogate the proton-motive force) readily confer tolerance to aminoglycosides (4).

### Strong evidence supports a role of TA modules in persister formation

Though we conclude that no evidence remains to support a role of the set of ten mRNA endonuclease toxin TA modules in persister formation of *E. coli* K-12 in unstressed cells, we continue to favor the hypothesis that at least some of these elusive genetic elements act as phenotypic switches into dormancy, a view largely shared by others in the field (1, 2, 31, 35). Others have confirmed important roles of different sets of TA modules in persister formation of *E. coli* K-12 and related organisms like uropathogenic *E. coli* and *Salmonella* (3, 4, 17, 36). With view to mRNA endonuclease TA modules, phenotypes of even single TA module deletions in persistence under specific conditions have repeatedly been described in the field such that, e.g., *yafQ* mutants of *E. coli* K-12 were found to be specifically defective in persister formation inside bacterial biofilms (37–39). Consistently, it is getting more and more clear that different TA modules in *E. coli* and beyond may respond to different upstream signaling (5, 40). We therefore argue that the role of mRNA endonuclease TA modules in persister formation of *E. coli* K-12 and a possible link to Lon deserve further investigation, even though our previously reported phenotypes with TA modules don’t hold true in the absence of φ80.

### Raising awareness of bacteriophage φ80

Beyond the confusion created by our erroneous interpretation of results obtained with φ80-infected strains, we are very worried that we have sent these strains to many research groups. Though it has been one of the favorite model phages in the early days of molecular biology (28), φ80 is now notorious as a laboratory contaminant. This bacteriophage combines “opportunistic induction” (i.e., a high frequency of spontaneous lytic development) with “stealth infectivity”, i.e., the comparably low infectivity of particles at 37°C and a relatively high preference for lysogeny (29, 30). Contamination of strains with φ80 can therefore go unnoticed for a long time – *E. coli* K-12 *Δ5TA* the first strain of the TA module deletion series carrying φ80, was first published by S. K. Christensen, et al. (41). However, elusive phenotypes such as a reduced efficiency of P1vir transductions can be strong hints of φ80 contaminations (29). Due to the inability of φ80 to adsorb to stationary phase cells, overnight cultures of lysogens regularly build up very high titers without any apparent sign of lysis (see also Figure 5C), facilitating contamination. Additionally, φ80 readily spreads through P1*vir* transductions from infected strains (29). We can therefore only suggest our colleagues to take the danger of φ80 contamination serious and use simple PCR tests like the one shown in Figure 4B to unravel if enigmatic phenotypes may be caused by φ80 infection. Of note, we have tested a random sample of (published) *E. coli* strains that we had obtained from different laboratories at different times and found several independent cases of φ80 infection.

### A refined methodology to study bacterial persister formation

Beyond the effects of φ80, we showed that the use of intermediate concentrations of fluoroquinolones (like 0.5-1 μg/ml of ciprofloxacin) for persister assays with *E. coli* is inappropriate, because - in full congruency with the results of E. L. Sandvik, et al. (25) for *S. aureus* – *ca*. one log of bacterial killing at these concentrations is not directly caused by the antibiotic but rather by the induction of resident prophages (Figure 2). It can therefore not be determined with certainty if phenotypes observed with experimental interventions or mutant strains are caused by differences in persister formation / survival or by effects on prophage induction. Future studies should therefore use higher concentrations of these antibiotics that inhibit prophage development, e.g., 10 μg/ml of ciprofloxacin.

Our results further suggest that commonly used protocols to study “exponential phase” persister formation are greatly affected by the carryover of dormant cells and / or molecules from overnight cultures (Figures 1 and 2). This effect may mask actual exponential phase persister formation until the absolute level of newly formed persister cells overcomes the level of inoculated dormant cells. It seems likely to us that this phenomenon may have contributed to the inconsistencies of results published by different laboratories, because persister assays with antibiotic treatment at the same cell density can give very different results depending on the amount of initial inoculum (Figure 1). Future studies should consider this effect when designing experiments and possibly, e.g., include controls of bacteria that are in balanced growth, i.e., have completed a sufficient amount of divisions since their inoculation from stationary phase so that effects of carryover can be excluded (42).

In this study we regularly used a very laborious setup of antibiotic killing assays that followed the levels of antibiotic-tolerant cfu/ml throughout the different growth phase of *E. coli* (see Figure 2A). Similar experiments have already been performed previously, e.g., in the seminal study by I. Keren, et al. (43). Though a number of experimental parameters differed between this study and our current work, there is clear agreement that β-lactam and fluoroquinolone tolerance is similar in early growth phases after inoculation and later increases strongly until it reaches (for β-lactams) full tolerance when bacteria cease growth (Figures 1 and 2; compare the results of I. Keren, et al. (43)). It seems likely that the stable level of tolerant cells after inoculation may reflect a long lag phase of dormant cells carried over from stationary phase, and others have indeed found growth lag to be the most important aspect of antibiotic tolerance for bacteria that are inoculated from stationary phase directly into fresh medium with antibiotics (44). It has therefore already been reasonably argued in the field that future studies should follow sufficiently elaborate methodologies in order to unravel the complexity of various antibiotic-tolerant subpopulations in bacterial cultures (16, 19, 21). We have also observed very interesting and highly divergent dynamics of antibiotic tolerance of *E. coli* K-12 to different antibiotics and in different media (Figures 1 and 2) that could not be explored in this study. However, it seems a promising field for future studies to compare the appearance and disappearance of tolerant cells for different antibiotics and try to uncover which external and intrinsic factors control antibiotic tolerance.

## Materials and Methods

### Bacterial strains and their construction

*E. coli* mutants were routinely constructed using recombineering with expression of the lambda red recombination genes from plasmid pWRG99 (45, 46). Clean deletions were constructed in a two-step procedure that first replaced the target gene(s) with the double-selectable cassette of template plasmid pWRG100, conferring chloramphenicol resistance for positive selection and carrying an I-SceI site for negative selection upon expression of the I-SceI endonuclease from pWRG99 (46). The double-selectable cassette was then removed through recombineering with a pair of annealed 80mer oligonucleotides spanning the desired deletion site with 40 basepair homologies on each side. All bacterial strains used in this study are listed in Table S1. The nucleotide sequences of all oligonucleotide primers are listed in Table S2.

*E. coli* K-12 MG1655 *relA::FRT spoT::cat(207)*, also known as PDC47, was constructed in two steps. The *relA* gene was deleted by P1vir transduction of the *relA::kanR* allele of the corresponding deletion strain of the KEIO collection and subsequent recombination of the flanking FRT sites using pCP20 (45, 47). In a second step, the *spoT::cat(207)* allele was transduced from *E. coli*CF1693 (48).

*E. coli* K-12 MG1655 *Δppkx*, also known as AHK062, was constructed by two-step recombineering as described with amplification of the double-selectable cassette using prAH1469/prAH1470 and 80mer oligonucleotides prAH1491/prAH1492.

*E. coli* K-12 MG1655 *sulA::FRT Δlon*, also known as AHK173, was constructed from *E. coli* K-12 MG1655 *sulA::kanR* of E. Maisonneuve, et al. (8) in two steps. We first removed the kanamycin resistance cassette by recombination of the flanking FRT sites using pCP20 (45) and then deleted *lon* by two-step recombineering with a double-selectable cassette amplified using prAH1465/prAH1466 followed by recombineering with 80mer oligonucleotides prAH1493/prAH1494.

*E. coli* K-12 MG1655 *Δ10’TA*, i.e., *E. coli* K-12 MG1655 *ΔhicAB::FRT ΔmqsR::FRT ΔyafO:::FRT* Δ*yhaV::FRT*Δ*higB::FRT ΔyefM-yoeB ΔdinJ-yafQ ΔrelBE ΔchpBS ΔmazF* (also known as AHK250) was constructed by sequential deletion of the remaining five TA modules / TA module toxins on the basis of *E. coli* K-12 MG1655 *A5’TA* of E. Maisonneuve, et al. (9) that has genotype *ΔhicAB::FRT ΔmqsR::FRT ΔyafO:::FRT* Δ*yhaV::FRT*Δ*higB::FRT*. This strain had been created by deleting five mRNA interferase toxins in reverse order to the TA module deletions of *Δ10TA.* Of note, *Δ5’TA* (unlike *Δ10TA* or *Δ5TA*) does not carry a lambda or φ80 prophage and does not show any difference to the parental wildtype in bacterial persister formation (Figure S4). The initial study by E. Maisonneuve, et al. (9) had described a substantial defect of the *Δ5’TA* strain in bacterial persistence, but we were unable to reproduce this phenotype (Figure S4), suggesting that the original work may have erroneously mixed up bacterial strains or inadvertently infected the clone used for experimentation with φ80. We deleted the *yefM-yoeB* locus in *Δ5′TA* using a double-selectable cassette amplified with prAH1633/prAH1652 and the 80mer oligonucleotide pair prAH1653/prAH1654. The *dinJ-yafQ* TA module was deleted with a double-selectable cassette amplified using prAH1627/prAH1649 and subsequent recombineering of 80mer oligonucleotides prAH1659/prAH1660. Subsequently, the *relBE* module was deleted using a double-selectable cassette amplified with primers prAH1631/prAH1651 and the 80mer oligonucleotides prAH1655/prAH1656. We deleted the *chpBS* locus with a double-selectable cassette amplified using prAH1648/prAH1626 followed by recombineering of 80mer oligonucleotides prAH1661/prAH1662. Finally, the *mazF* toxin gene was deleted using a double-selectable cassette amplified with prAH1629/prAH1650 and the 80mer oligonucleotides prAH1657/prAH1658. We confirmed the successful introduction of all deletions in *Δ10’TA* and also parental *Δ5’TA* using diagnostic PCRs over the TA loci.

*E. coli* K-12 MG1655 *Φ80(+)* was generated by lysogenization of the parental wildtype with φ80 of strain *Δ10TA attB(+)*. In short, culture supernatant of *E. coli Δ10TA attB(+)* was prepared as described for plaque assays below and 100 μl were added to a culture of *E. coli* K-12 MG1655 at OD600 of ca. 0.2. The culture was agitated at 30°C until lysis occurred and survivors were plated on LB agar plates. After overnight incubation at 37°C, single colonies were tested for a K-12 MG1655 wildtype chromosomal background (by PCR on TA module loci) and carriage of only wildtype φ80 (by sensitivity to lambda *cl*_*b221*_).

### Curing of the lambda prophage from *E. coli* K-12 *Δ10TA*

The original *E. coli Δ10TA* strain of E. Maisonneuve, et al. (9) was cured of its lambda prophage via a two-step P1wr transduction procedure. In short, the lambda prophage was first replaced with a temperature-sensitive λ.RED variant, and this prophage was subsequently cured by P1vir transduction of a native *gal-bio* region with an unoccupied *attB* integration site from the *E. coli* K-12 MG1655 wildtype strain.

Strain HME71 of J. A. Sawitzke, et al. (49) carries a temperature-sensitive λRED prophage and was cured for its *Δ(srlA-recA)301::Tn*10 insertion by P1*vir* transduction of the native locus from the *E. coli* K-12 MG1655 wildtype strain in order to restore tetracycline sensitivity by removal of the resistance marker encoded on the Tn10 transposon. Transduction of the wildtype locus was selected as sorbitol prototrophy (Srl^+^) and verified by screening for a Rec^+^ phenotype and tetracycline sensitivity. Subsequently, the resulting strain (MAS889) was transduced back to tetracycline resistance with a P1*vir* lysate raised on strain MAS242 that carries a mini-Tn10 inserted close to *galE* near the lambda attachment site *attB,* thus establishing a tetracycline resistance marker that is genetically linked to λ.RED. Transductant colonies were screened for temperature sensitivity (i.e., presence of λRED). A P1*vir* lysate of the resulting strain, MAS902, was used to transduce tetracycline resistance into *E. coli Δ10TA* in order to replace the resident lambda prophage with λRED. Successful replacement was verified by screening for temperature sensitivity. Subsequently, the wildtype *gal-bio* region with an unoccupied *attB* site was transduced from *E. coli* K-12 MG155 by selection for galactose and biotin prototrophy (Gal^+^ Bio^+^) and loss of temperature sensitivity, creating strain *E. coli Δ10TA attB(+).* Loss of tetracycline resistance was verified and successful curing of lambda prophages was confirmed by the detection of an unoccupied *attB* site using an overspanning PCR with primer pair prMASgal/prMASbio of K. Baek, et al. (50). Analogous efforts to cure φ80 infections from lysogens have remained unsuccessful, as reported earlier by others (29).

### Preparation of culture media

Luria-Bertani (LB) broth was prepared by dissolving 10 g of tryptone (cat#LP0042, Oxoid), 5 g of yeast extract (cat#LP0021, Oxoid), and 10 g of sodium chloride (cat#27810.364, VWR Chemicals) per liter of Milli-Q H_2_O and sterilized by autoclavation. M9 medium was prepared as “M9 minimal medium (standard)” of Cold Spring Harbor Protocols (51) with modifications as 1× M9 salts (stock solution prepared from 5× M9 salts, cat#M6030 Sigma-Aldrich, supplemented with 50 μl of a 10 mg/ml FeSO_4_ solution (cat#F8048, Sigma)), 0.4% w/v Bacto casamino acids (cat#223020 BD Biosciences; from 20% w/v sterile-filtered stock), 0.4% w/v D-glucose (cat#101176K, VWR Chemicals, from 40% w/v stock), 2 mM MgSO_4_,(cat#25.165.292, VWR Chemicals) 1 μg/ml thiamine (cat#T1270, Sigma), 100 mM CaCl_2_,(cat#26.764.298, VWR Chemicals). For the (p)ppGpp-deficient strain *E. coli* K-12 MG1655 *relA::FRT spoT::cat(207)*, casamino acids in the M9 medium were replaced with 400 μg/ml of L-serine and 40 μg/ml of all other amino acids (all ≥98% or ≥98% purity, Sigma) to support reasonable growth of this delicate strain as described by K. Potrykus, et al. (32).

### Persister assays

The presence of persister cells in bacterial cultures is usually detected using biphasic kill curves in which the addition of a bactericidal antibiotic is first followed by rapid killing of regular cells and then a second, slower phase in which persister cells are killed (1, 2). Persister cells can then be quantified by comparing the levels of survivors obtained with, e.g., different mutant strains after a given time of antibiotic treatment. Experimental procedures similar to the one used here had also been described in our previous work and numerous studies by others in the field (5, 8, 9, 11). These studies showed that antibiotic treatment in LB medium and different minimal media at exponentially growing bacteria of ca. 1 - 5 × 10^8^ cfu / ml results in biphasic killing with merely persisters surviving after 5 hours. Similarly, we demonstrate biphasic killing under these conditions for the M9 minimal medium that we used throughout this work (Figure S2).

Overnight cultures were inoculated from single colonies into 3 ml of LB or M9 medium and grown for ca. 16 hours in plastic culture tubes (cat#62.515.006, Sarstedt; lid arrested with tape at the 13-ml mark with its lower end to ensure uniform aeration of replicates). For persister assays based on a single growth time-point as described in Figure 1A, overnight cultures were diluted back 1:100 into LB or M9 medium in Erlenmeyer flasks and agitated at 37°C in a waterbath shaker until they were in mid-exponential growth phase (ca. 1 - 5 × 10^8^ cfu / ml; reached in our setup after ca. 2 – 2.5 hours in LB medium or 2.5 – 3 hours in M9 medium). At this point cultures were treated with lethal concentrations of different antibiotics (100 μg/ml ampicillin, 1 or 10 μg/ml ciprofloxacin, or 7.5 μg/ml gentamicin) for 5 hours in plastic culture tubes under rigorous agitation. In parallel, cfu/ml of the cultures were determined by plating serial dilutions on LB agar plates. After antibiotic treatment, bacterial pellets of 1.5 ml samples were washed once in 1 ml of sterile phosphate buffered saline (PBS), resuspended in 100 of sterile PBS, and serially diluted in sterile PBS. 10 samples were spotted on LB agar plates to quantify antibiotic-tolerant survivors. Agar plates were incubated at 37°C for at least 24 h and cfu / ml were determined from spots containing 10 – 100 bacterial colonies. The fraction of persister cells was calculated as the ratio of cfu/ml after and before antibiotic treatment.

In order to study the dynamics of antibiotic-tolerant cells throughout the different growth phases of *E. coli* (as outlined in Figure 2A), we performed an experiment as described above but varied the time between subculturing and antibiotic treatment from 0 hours (i.e., direct inoculation into fresh medium containing antibiotic) to 8 hours. For experiments with the (p)ppGpp-deficient strain *E. coli* K-12 MG1655 *relA::FRT spoT::cat(207)*, the M9 growth medium was supplemented with an amino acid mixture instead of casamino acids (see above) and the experiment was extended to 10 hours. Though the *relA spoT* mutant and *E. coli* K-12 wildtype have roughly the same growth rate in this medium, the final cfu/ml of the (p)ppGpp-deficient mutant is significantly lower which regularly results in experimental artifacts due to the appearance of suppressor mutants during overnight cultures (32). We therefore set up overnight cultures in LB medium (where this effect is much less pronounced) and adjusted the inoculum of both mutant and wildtype ad ca. 10^6^ cfu/ml in order to account for the roughly 3-fold higher cfu/ml of wildtype overnight cultures.

### Genome sequencing and analysis

Genomic DNA of *E. coli* K-12 MG1655 wildtype as well as its *Δ5TA*, *Δ8TA*, and *Δ10TA attB(+)* derivatives was isolated by phenol chloroform extraction and sequenced by GATC Biotech on the Illumina Hiseq 2500 sequencing platform to a read length of 51 basepairs. Between five and ten million paired end reads were obtained for each strain, resulting in an average 200-fold sequencing depth. Sequencing reads were assembled using the Velvet assembler (52) in both *de novo* and referenced modes, where the previously published genome of *E. coli* K-12 MG1655 wildtype (accession U00096) served as a template for the latter. Suspected prophage infection was detected by assembling the raw contigs (around 20 per strain) to the published *E. coli* K-12 MG1655 genome sequence (accession U00096) and examining those not matching the reference by BLAST searches.

### Prophage detection by PCR

Prophage carriage was detected by PCR with oligonucleotide primer pairs that amplified the two junctions between the *E. coli* K-12 chromosome and the prophage in the *gal-bio* region (lambda) and at the *ycil* locus (φ80). Bacteriophage lambda was detected using primer pairs prMASgal/prMASint (*gal-int*) and prMASb2/prMASbio (*b2-bio*) that have been used previously for this purpose (50). Bacteriophage φ80 was detected using primer pairs prAH1506/prAH1499 (*kch-int*) and prAH1500/prAH1521 (*pinL-tonB*). The nucleotide sequences of all oligonucleotide primers are listed in Table S2.

### Phage techniques

Our phage work followed standard techniques that had already been used to study φ80 infections by others (29, 30). Plaque assays were performed by spotting lytic phages or correspondingly treated supernatants of *E. coli* cultures on LB plates overlaid with a top agar containing *E. coli* strains as indicated. Lytic phage stocks were produced by diffusion of phage particles into SMG buffer (0.1M NaCl, 10 mM MgSO_4_, 0.05M Tris pH7.5, 0.01% gelatin in Milli-Q H_2_O) from top agar plates with confluent plaques on *E. coli* K-12 MG1655. Supernatants of overnight cultures of suspected φ80 lysogens or control strains were separated from bacterial material by centrifugation and then diluted 1:10 into SMG buffer containing 10% chloroform. For plaque assays, LB agar plates in square format (12 cm × 12 cm) were overlaid with 7ml of LB top agar (0.7% agarose) containing 100 μl of an overnight culture of indicated *E. coli* strains. After the top agar had solidified, 5 of phage of supernatant stocks were spotted on the top. Plates were incubated at 30°C (to enable efficient plaque formation by φ80 (29)) overnight until plaque formation was visible. Sensitivity of bacterial strains to lambda *cl*_*b221*_ was assayed by evaluation the growth of bacterial streaks that crossed lines of phage stock on LB agar plates supplemented with 20 mM MgSO_4_ and 5 mM CaCl_2_. The ability of various strains to grow after contact with lambda *cl*_*b221*_ was evaluated visually after overnight incubation of agar plates at 37°C.

### Quantification and statistical analysis

Experiments were usually analyzed by calculating mean and standard deviation of at least three biological replicates. Detailed information for each experiment is provided in the figure legends.

## Acknowledgments

The authors thank Etienne Maisonneuve for the preparation of *E. coli* DNA and Shiraz A. Shah for the processing of DNA sequences. They are particularly grateful to Sine Lo Svenningsen for *E. coli* JMT1, Patricia Dominguez-Cuevas for *E. coli* PDC47, and Anders Løbner-Olesen for *E. coli* Δ9CP. Steen Pedersen and Stanley Brown are acknowledged for helpful discussions about φ80.

## Supplemental Figure Legends

**Figure S1. The experiment shown in Figures 1B and 1C performed with a different batch of LB medium.** We performed the experiment shown in Figures 1B and 1C with a different batch of LB medium and compared cfu/ml (A) as well as the fraction of antibiotic-tolerant cells (B) at treatment with ciprofloxacin ad 1 μg/ml. Note that the growth of regular bacteria is not different between batch #1 (shown in Figure 1) and batch #2 (shown here), but that the dynamics of persister levels differ at early time-points.

**Figure S2. Biphasic killing of exponentially growing *E. coli* K-12 MG1655 in M9 medium.** Bacteria were cultured as described for the experiments shown in Figure 2 and treated with 100 μg/ml ampicillin, 1 μg/ml ciprofloxacin, or 7.5 μg/ml gentamicin. Samples were withdrawn at different time-points for 5 hours and the number of surviving cfu/ml was plotted against the duration of treatment. Note that *E. coli* K-12 displays biphasic killing when challenged with any of the three antibiotics while growing exponentially in M9 medium, suggesting that tolerance assays at this growth phase report bacterial persister cells. Data points represent the mean of at least three independent experiments and error bars indicate the standard deviation.

**Figure S3. No phenotype of *sulA::FRT* in antibiotic tolerance.** (A,B) As a control for the experiments with *E. coli* K-12 MG1655 *sulA::FRT Δlon* strain shown in Figure 7, we studied the dynamics of antibiotic tolerance of the parental *sulA::FRT* mutant and plotted cfu/ml (A) as well as the fraction of antibiotic-tolerant cells (B). As expected from the literature, the *sulA* mutant strain showed no defect in antibiotic tolerance (see the study by A. Theodore, K. Lewis, and M. Vulic, Genetics 195:1265-1276, 2013, doi:10.1534/genetics.113.152306). Data points show the mean of three independent experiments and error bars represent the standard deviation.

**Figure S4. The *Δ5’TA* strain displays no phenotype in bacterial persistence.** Exponentially growing cultures of *E. coli* K-12 MG1655, the *Δ5’TA* strain, or the *Δ10TA attB(+)* strain were challenged with 1 μg/ml ciprofloxacin for 5 hours in LB medium and the fraction of surviving persister cells was determined. Data points represent the mean of three independent experiments and error bars indicate the standard deviation.

